# Functional diversity of isoprenoidal lipids in *Methylobacterium extorquens* PA1

**DOI:** 10.1101/2020.12.21.423902

**Authors:** Sandra Rizk, Petra Henke, Carlos Santana-Molina, Gesa Martens, Marén Gnädig, Damien P Devos, Meina Neumann-Schaal, James P Saenz

**Author notes:** Address correspondence to James P Saenz.

## Abstract

Hopanoids and carotenoids are two of the major isoprenoid-derived lipid classes in prokaryotes that have been proposed to have similar membrane ordering properties as sterols. *Methylobacterium extorquens* contains hopanoids and carotenoids in their outer membrane, making them an ideal system to investigate whether isoprenoid lipids play a complementary role in outer membrane ordering and cellular fitness. By genetically knocking out *hpnE*, and *crtB* we disrupted the production of squalene, and phytoene in *Methylobacterium extorquens* PA1, which are the presumed precursors for hopanoids and carotenoids, respectively. Deletion of *hpnE* unexpectedly revealed that carotenoid biosynthesis utilizes squalene as a precursor resulting in a pigmentation with a C_30_ backbone, rather than the previously predicted C_40_ phytoene-derived pathway. We demonstrate that hopanoids but not carotenoids are essential for growth at high temperature. However, disruption of either carotenoid or hopanoid synthesis leads to opposing effects on outer membrane lipid packing. These observations show that hopanoids and carotenoids may serve complementary biophysical roles in the outer membrane. Phylogenetic analysis suggests that *M. extorquens* may have acquired the C_30_ pathway through lateral gene transfer with Planctomycetes. This suggests that the C_30_ carotenoid pathway may have provided an evolutionary advantage to *M. extorquens*.

**Importance:** All cells have a membrane that delineates the boundary between life and its environment. To function properly, membranes must maintain a delicate balance of physical and chemical properties. Lipids play a crucial role in tuning membrane properties. In eukaryotic organisms from yeast to mammals, sterols are essential for assembling a cell surface membrane that can support life. However, bacteria generally do not make sterols, so how do they solve this problem? Hopanoids and carotenoids are two major bacterial lipids, that are proposed as sterol surrogates. In this study we explore the bacterium *M. extorquens* for studying the role of hopanoids and carotenoids in surface membrane properties and cellular growth. Our findings suggest that hopanoids and carotenoids may serve complementary roles balancing outer membrane properties, and provide a foundation for elucidating the principles of surface membrane adaptation.

## Introduction

Microorganisms can withstand a diversity of environmental stresses ranging from extreme temperatures to the immune defenses of multicellular organisms. The cellular surface membrane serves as a first line of defense against environmental perturbations and the membrane’s lipid composition is critical for stress resistance. On the one hand the membrane must be robust enough to withstand chemical and physical challenges. On the other hand, the membrane must be fluid enough to support bioactivity. In eukaryotic organisms such as yeast, sterols play a crucial role in achieving a fluid yet mechanically robust cell surface membrane^1^. However, bacteria generally do not synthesize sterols with very few exceptions^2,3^.

The absence of sterols from most prokaryotes suggests that alternate lipids may serve analogous roles in surface membranes. All three domains of life possess isoprenoid synthesis pathways derived from a common C_5_ isoprene building block which give rise to a broad suite of diverse lipid classes including sterols, but also carotenoids and hopanoids, and the majority of archaeal lipids. Because of their structural similarities that are derived from a common C_5_ isoprene building block, resulting in rigid and often semi-planar structures, isoprenoid-derived lipids may share certain biophysical features in membranes^3^. However, the mechanism and exact influence of isoprenoid lipids on prokaryotic membrane properties and cellular fitness remains relatively unexplored.

There is increasing evidence pointing to the role of bacterial isoprenoid-derived lipids such as hopanoids and carotenoids in membrane stabilization in bacteria^4^. Hopanoids have been shown to order outer membrane lipids by interacting with lipid A in a similar manner to that exhibited by cholesterol and sphingolipids in eukaryotes^5–7^. Whereas, carotenoids (ß-carotene and zeaxanthin) have been shown using molecular dynamics (MD) simulations to have a condensing effect similar to that of cholesterol on phospholipids^8^. Physiologically, there is evidence that hopanoids are important for growth at higher temperatures^9–13^, whereas carotenoids have been linked to cold acclimation in some bacteria^14–16^. These contrasting phenotypes for temperature acclimation suggest that hopanoids and carotenoids may serve complementary roles in modulating membrane properties. Taken together, these observations suggest functional similarities between sterols and bacterial isoprenoid lipids. However, the extent to which carotenoids and hopanoids have similar biophysical properties and functions in biomembranes is not known and has not been systematically explored in a living model system.

*Methylobacterium extorquens* is a Gram-negative bacterium with a well characterized genome and a simple lipidome^17^, that produces both hopanoids and carotenoids. This makes it an attractive model organism for studying the global phenotypes of disrupting the two pathways. *M. extorquens* has been shown to overproduce carotenoids when the gene squalene hopene cyclase (*shc*) is knocked out^18^. However, whether there is cross-talk between the two pathways is not known. In this study we have genetically disrupted the biosynthetic pathways of the two main isoprenoid lipid precursors; squalene (precursor for hopanoids) and phytoene (precursor for carotenoids), thus confirming the function of the gene *hpnE* in *M. extorquens*, additionally, we show that even though the genome of *M. extorquens* has the genes for the C_40_-carotenoids biosynthetic pathway, the pigmentation has a C_30_-based backbone that is squalene derived. We demonstrate the importance of hopanoids for growth at high temperatures, implicating its role in membrane temperature adaptation. By measuring lipid packing in the outer membrane we show that deletion of carotenoids and hopanoids results in opposing changes in membrane properties, raising the possibility that hopanoids and carotenoids collectively serve diverse but complementary roles in maintaining membrane properties. Finally, we propose that the genes for the C_30_ squalene derived pathway were acquired through lateral gene transfer (LGT), suggesting that producing C_30_ carotenoids provides a selective evolutionary advantage over C_40_ carotenoids in this organism.

## Results

### Confirming the function of the genes *hpnE* and *crtB* in *M. extorquens* PA1

We first knocked out the isoprenoid precursors phytoene and squalene by deleting the genes phytoene synthase (*crtB*) and hydroxysqualene oxidoreductase (*hpnE*). We measured the absorbance spectra of lipids extracted from the strains WT, Δ*crtB* and Δ*hpnE* as a read out for carotenoid pigmentation. The Δ*crtB* mutant strain showed no loss in pigmentation compared to the WT (**Figure 1A**), whereas the Δ*hpnE* mutant strain was non-pigmented (**Figure 1B**). Indeed, LC-MS analysis revealed that the Δ*hpnE* strain no longer produced detectable amounts of diplopterol (hopanoids) (**Table 1**), and that it accumulated hydroxysqualene which is the precursor of squalene biosynthesis (**Table 1**)^19^. Combined, these observations led us to investigate whether carotenoid biosynthesis in *M. extorquens* is derived from squalene^20^ rather than phytoene.

**Figure 1.**
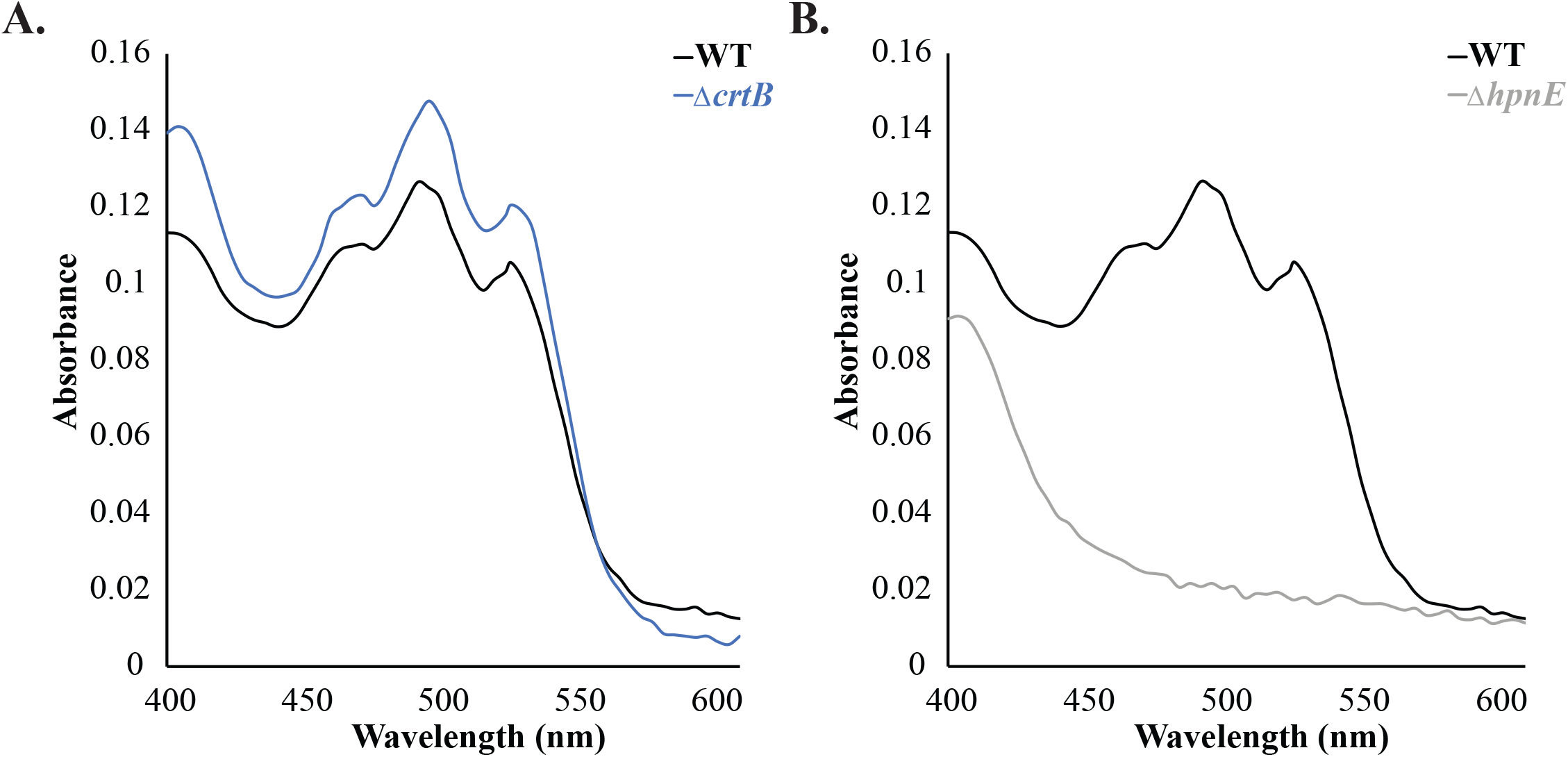
Confirmed the function of the genes *hpnE* and *crtB* in *M. extorquens* PA1. **A**. Absorbance spectrum of lipids extracted from Δ*crtB*, and **B**. Δ*hpnE* strains normalized to cell mass.

**Table 1:** Semiquantitative measurement of integrated peak area obtained from LC-MS analysis, normalized to 10 mg wet weight of cells performed on lipid extracts from the strains WT, Δ*shc*, Δ*hpnE*

| Compound | WT | $\Delta shc$ | $\Delta hpnE$ |
| --- | --- | --- | --- |
| Hydroxysqualene | n.d | n.d | 84.8±10.1 |
| Diplopterol | 19±1.4 | n.d | n.d |
| Squalene | n.d | 318.4±2.8 | n.d |

### Carotenoids are derived from the C_30_ pathway in *M. extorquens*

In order to confirm that carotenoid biosynthesis uses squalene as a precursor in *M. extorquens*, we knocked out the genes in the C_30_ pathway (*crtN*, *crtP*)^20^, analyzed the absorbance spectra of the lipids extracted from different mutant strains, and performed LC-MS on the pigments to determine the chemical composition of their carbon backbone. We observed that knocking out *crtN*, and *crtP* resulted in loss of pigmentation (**Figure 2A, B**), whereas, knocking out *crtB* did not (**Figure 1A**), hence, we amended the carotenoids biosynthetic pathway in *M. extorquens* PA1 (**Figure 2C).** Moreover, LC-MS analysis confirmed that carotenoids detected in *M. extorquens* all have a C_30_ backbone, and no C_40_ backbone-based carotenoids were detected confirming its squalene origin (**Figure S1**). The Δ*shc* mutant exhibited more pigmentation due to increased carotenoid production (**Figure 2D**) in agreement with the findings of Bradley et al.^18^. We observed an accumulation of squalene in the Δ*shc* strain as detected by LC-MS, which could explain the increase in synthesis of carotenoids (**Table 1**, **Figure 2D**). The deletion of genes in the proposed squalene-derived C_30_ carotenoid pathway produced non-pigmented mutant strains, where the phenotype was eliminated by gene complementation on an inducible plasmid (**Figure S2**). Whereas knocking out genes in the C_40_ carotenoids biosynthesis pathway had no effect on pigmentation, LC-MS analysis confirmed the presence of a C_30_ backbone of the carotenoids pigment extracted from the WT, Δ*shc* and Δ*crtB* strains. These results suggest that C_40_ biosynthetic pathway was not active, or at least, it was not expressed at optimal growth conditions in *M. extorquens*.

**Figure 2.**
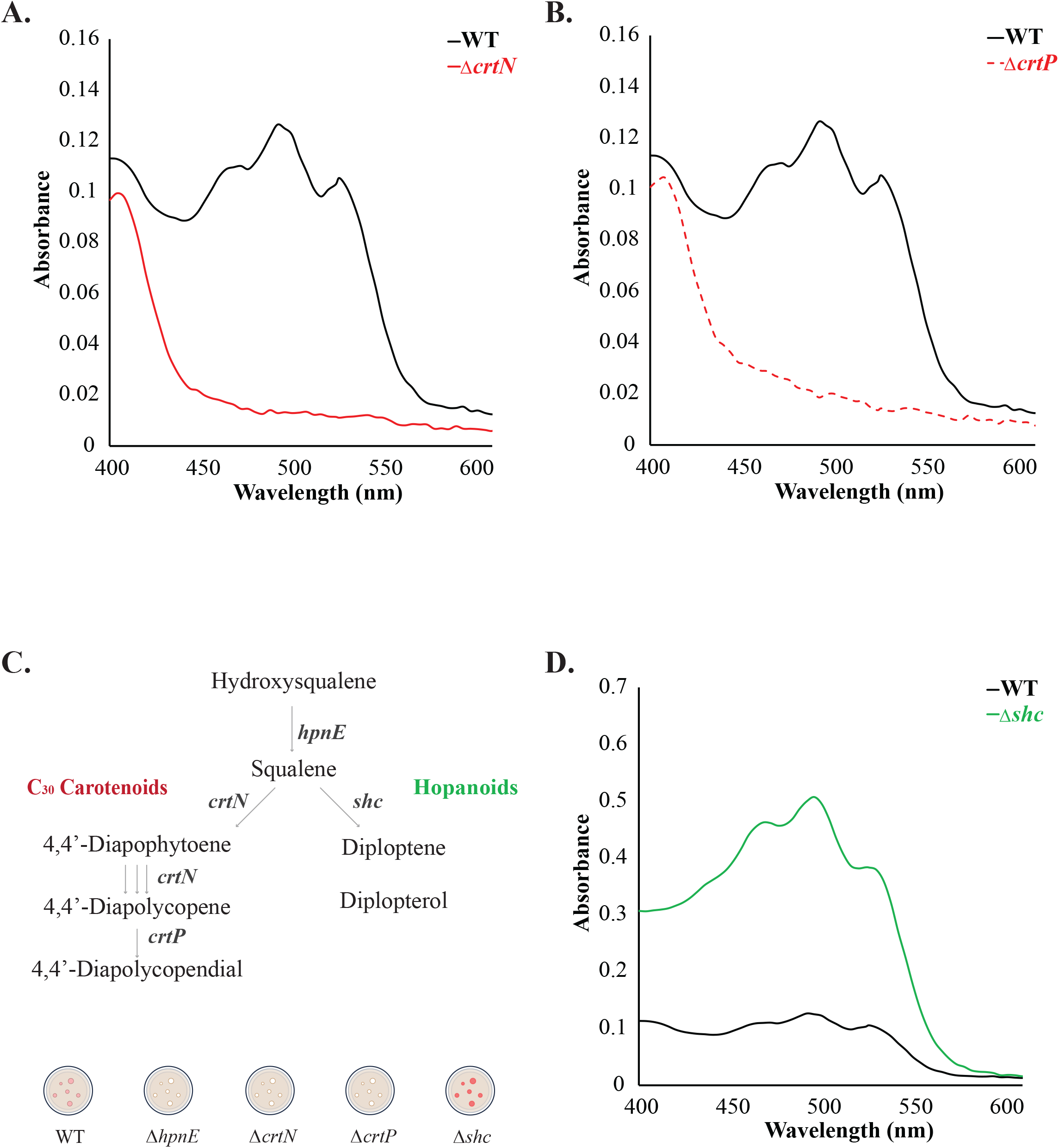
Identification of the genes of squalene derived C_30_ carotenoids biosynthetic pathway in *M. extorquens* PA1. **A.** Absorbance spectra normalized to cell mass of lipids extracted from mutant strains in the C_30_ pathway Δ*crtN*, and **B.** Δ*crtP* which resulted in loss of pigmentation. **C.** Amended carotenoids biosynthetic pathway upon knocking out genes in the C_30_ pathway thus confirmed their function due to observed loss in pigmentation. **D.** Absorbance spectra of lipids extracted from Δ*shc* mutant strain depicted an observed increase in pigmentation (normalized to cell mass).

### Growth phenotypes of isoprenoids mutants at different temperatures

In order to explore how hopanoids and carotenoids contribute to cellular growth and acclimation to environmental stresses, we investigated how disrupting the biosynthesis of hopanoids and carotenoids affected cellular growth at different temperatures. Temperature change is a key environmental stress that *M. extorquens* must withstand in its native habitat on plant leaves, where it can experience wide diurnal variations. We previously showed that temperature has one of the largest effects on lipidomic remodeling and growth rate, relative to other experimental parameters such as detergent and salt concentrations^17^. Here, we demonstrated that interrupting hopanoid biosynthesis (Δ*hpnE*, Δ*shc*) caused a growth impairment especially at temperatures higher than the optimum (30°C) (**Figure 3A**). Moreover, increased carotenoid production by the Δ*shc* strain did not rescue the growth phenotypes observed, on the contrary a more adverse effect was detected as compared to the Δ*hpnE* strain (**Figure 3A**). On the other hand, knocking out the C_30_ biosynthetic desaturases (*crtN*, *crtP*) did not have any effect on growth at different temperatures (**Figure 3B**). These results confirm the dependence of heat tolerance on hopanoids in *M. extorquens*. Whereas, carotenoids did not seem to play a crucial role in membrane’s temperature stabilization in this organism.

**Figure 3.**
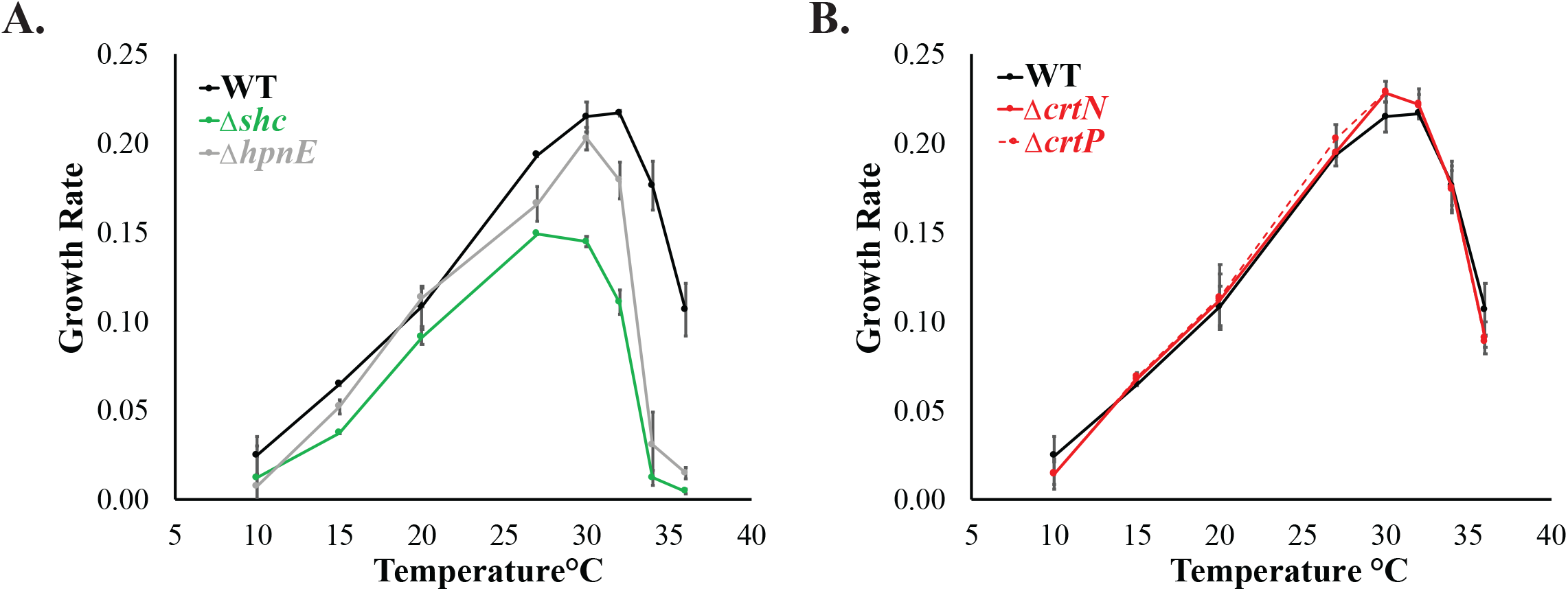
Effect of disrupted isoprenoid biosynthesis on growth at different temperatures. **A.** Hopanoid knockout strains comparison with WT, **B.** C_30_ carotenoids knockout strains comparison with WT.

### Growth impairment at high temperature in hopanoid knockout strains is associated with an increase in membrane fluidity that cells cannot compensate for

The high temperature growth impairment observed for mutant strains that cannot produce hopanoids implicated a membrane-induced defect. To investigate the membrane properties of different hopanoid knockout strains, we used the lipophilic dye Di-4 ANEPPDHQ (Di-4) which reports on lipid packing through the calculation of general polarization (GP), thus higher GP indicates more packed lipids^21^. Lipid packing is correlated with a number of key membrane properties including viscosity and bending rigidity, thereby providing a robust and sensitive readout of variations in the physical state^22,23^. It has been previously shown that Di-4 selectively labels the surface membrane, most likely due to its bulky polar headgroup which prevents flipping to the inner leaflet^24^. We therefore measured the lipid packing *in vivo* of the isoprenoid-lipid mutant strains: WT, Δ*crtN*, Δ*crtP*, Δ*shc* and Δ*hpnE*. Our first observation was that the GP values obtained for WT and Δ*shc* strains *in vivo* were comparable to the GP values reported by Sáenz et al. for the *in vitro* measurement on purified outer membranes^6^. Our findings showed that Δ*shc*, and Δ*hpnE* mutant strains had a much lower GP which indicated less lipid packing as compared to the WT strain even at their optimal growth temperature, whereas, Δ*crtN*, and Δ*crtP* strains had increased lipid packing compared to the WT strain (**Figure 4A**). These results imply that even in the native state, the outer membrane lipids of the hopanoid knockout strains were less packed as compared to the WT strain. In addition, the loss of carotenoids (Δ*crtN*, Δ*crtP*) seemed to slightly increase lipid packing. We then measured the GP at the maximum growth temperature of the hopanoid mutants (32°C), and we showed that there is no marked change in GP (ΔGP) for the WT strain as opposed to Δ*shc* and Δ*hpnE* strains (**Figure 4B**). We propose that cells preserve a certain range of parameters to maintain their vitality and ability to survive challenging environmental perturbations. Hence, the removal of hopanoids highly restricts their fitness at higher temperatures by compromising the cellular adaptability.

**Figure 4.**
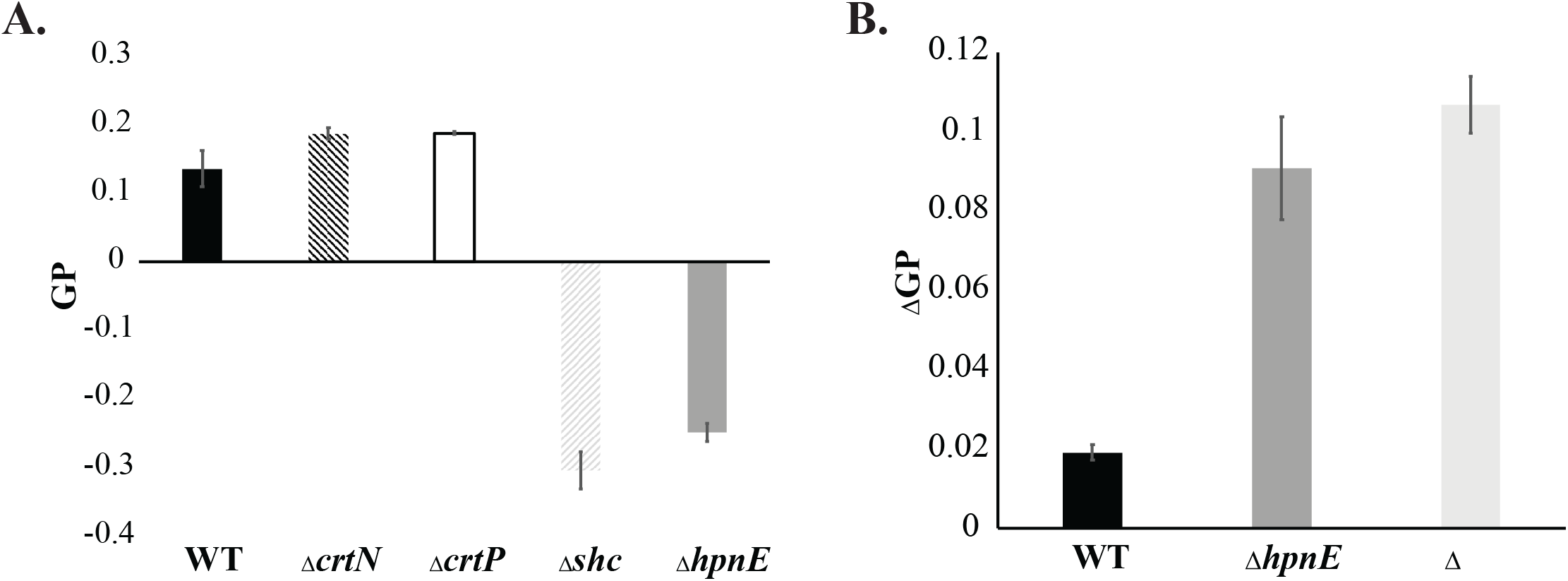
Effect of loss of membrane isoprenoids on lipid packing. **A.** Outer membrane general polarization (GP) as measured by Di-4-ANEPPDHQ for hopanoids knockout strains (Δ*shc*, Δ*hpnE*), and carotenoids knockout strains (Δ*crtN*, Δ*crtP*) at 27°C. **B.** Difference in GP of cells grown at 27°C and 32°C and the change in GP (ΔGP) reported.

### Phylogeny reveals co-occurrence of C_30_ and C_40_ pathway in *M. extorquens* and suggests C_30_ pathway was acquired through horizontal gene exchange

Carotenoid biosynthesis in *M. extorquens* has been previously hypothesized to stem from phytoene^25^. Nonetheless, the loss of pigmentation observed in the Δ*hpnE* mutant strain suggested that carotenoids are squalene derived (C_30_-based carotenoid backbones). Hence, we analyzed the distribution of both pathways in Proteobacteria (**Figure 5A**). Additionally, we performed the phylogeny of the FAD-dependent desaturases which are involved in the initial steps of C_40_ and C_30_ carotenoid biosynthesis (**Figure 5B**). We found in the *M. extorquens* genome the genes coding for the enzymes CrtB-CrtD-CrtI (for C_40_ carotenoids), CrtN-CrtP (for C_30_ carotenoids) and HpnCDE (for C_30_ squalene) (**Figure 5A**). The phylogeny of the respective CrtD-CrtI enzymes located *M. extorquens* sequences branching within the Alpha- and Gammaproteobacteria group (**Figure 5B**). The phylogeny of HpnCDE enzymes showed monophyly of Alpha- and Gammaproteobacteria^26^, but HpnCDE appeared more conserved than CrtB-CrtD-CrtI^26^ (**Figure 5A**). This monophyly of Alpha- and Gammaproteobacteria suggested that both squalene and C_40_ carotenoid biosynthesis were ancestral in Proteobacteria. By contrast, the C_30_ FAD-dependent desaturase enzymes CrtN and CrtP, displayed a more limited distribution in Alphaproteobacteria, particularly in Rhodospirillales, Rhizobiales, Acetobacterales, Azospirillales orders (taxonomic orders according to GTDB; **Figure 5A**). In addition, these sequences, including the *M. extorquens* ones, did not branch close to, nor monophyletically with, the Gammaproteobacteria. Instead, the respective alphaproteobacterial groups of CrtN and CrtP branched within the Planctomycetes (**Figure 5B**), suggesting lateral gene transfer (LGT) from this group (**Figure 5B**). Planctomycetes are a distant bacterial phylum that had recently been proposed to produce C_30_ carotenoids via squalene synthesis enzymes HpnCDE^26^. The similar topology between CrtN and CrtP branches (**Figure 5B**) suggested that these genes were transferred together i.e. in the same DNA fragment/locus. Therefore, unlike CrtI-CrtD or HpnCDE enzymes which indicated an ancestral feature of Proteobacteria, the C_30_ carotenoid pathway in some Alphaproteobacteria orders suggest that they originated later by LGT from Planctomycetes.

**Figure 5.**
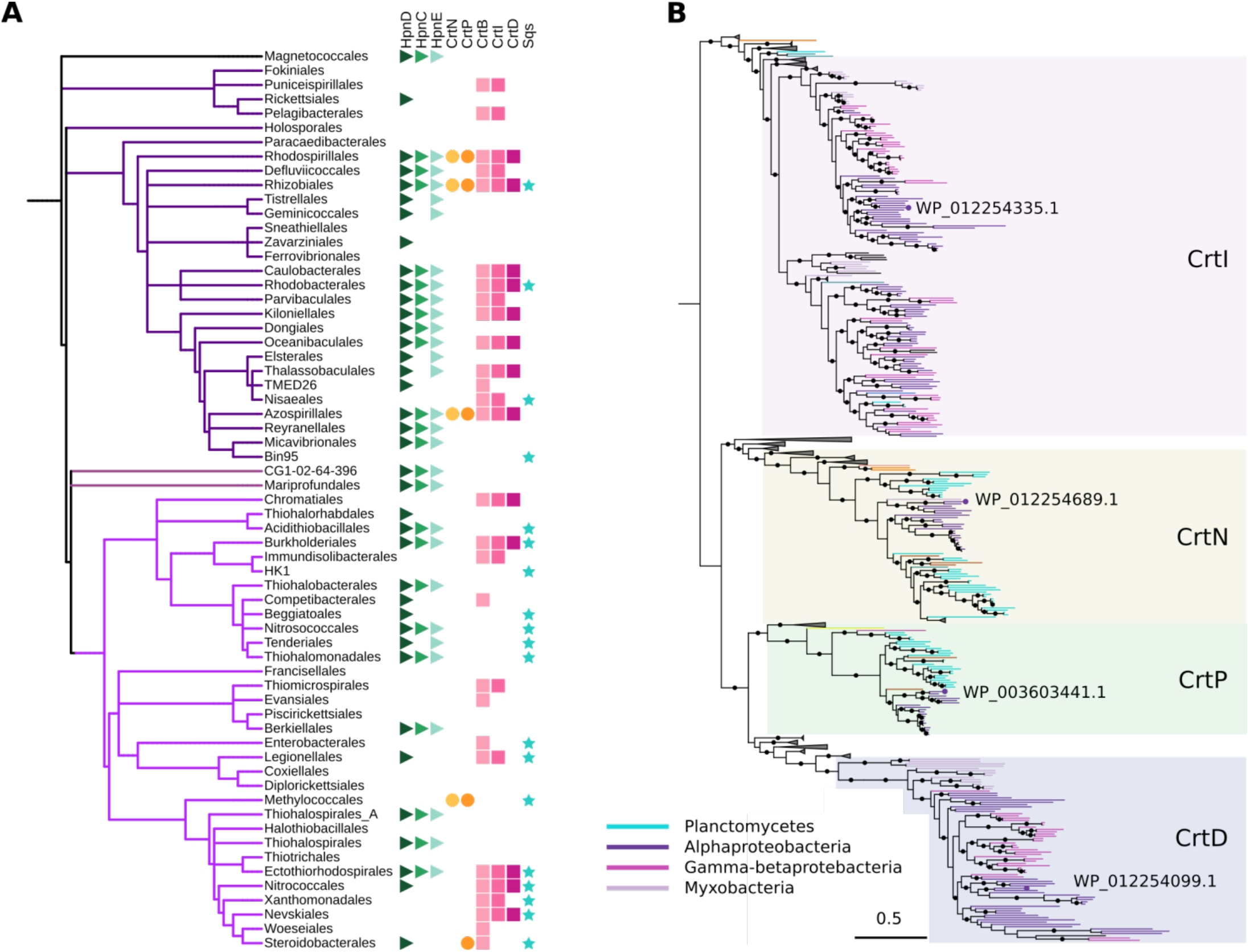
**A.** Phylogenetic profile of squalene and carotenoid related enzymes (Right) mapped onto a taxonomic tree of orders from Proteobacteria (Left). The tree was obtained from the Genome Taxonomy Database^46^ pruning the branches of interest. **B.** Phylogeny of the amino oxidases involved in carotenoid biosynthesis. The *M. extorquens*’ protein codes are shown. The branches are colored according to the taxonomy and other branches were collapsed to ease the visualization. The subfamilies are annotated according to the presence of characterized proteins. Black dots indicate bootstraps higher than 90.

## Discussion

In order to establish *M. extorquens* as a system to study the comparative role of carotenoids and hopanoids in determining membrane properties, we first determined how to perturb biosynthesis of the two pathways independently. Hopanoid biosynthesis in *M. extorquens* has been relatively well described^27^, however the squalene synthase was never formally identified or confirmed by a knockout strain. We identified and confirmed the function of *hpnE* as a key gene that would disrupt squalene synthesis^19^, thereby disrupting hopanoids biosynthesis and preventing squalene accumulation^28–30^. Under the assumption that carotenoids were derived from phytoene in *M. extorquens*^25^, we targeted a phytoene synthase gene *crtB*. However, surprisingly deletion of *crtB* showed no phenotype in pigmentation, whilst deletion of *hpnE* yielded non-pigmented mutant strains that also lacked hopanoids. These unexpected results revealed that carotenoid synthesis was derived from squalene rather than phytoene through a pathway that has recently been shown to produce C_30_ carotenoids^20^.

Having identified the genes required to independently disrupt the hopanoid and carotenoid pathways, we investigated the effects of varying temperature on cellular growth rates in strains deficient in either hopanoids or carotenoids. Hopanoids have been shown to modulate bacterial membrane properties in a manner analogous to eukaryotic sterols^5,6^. In *M. extorquens* deletion of hopanoid synthesis by deleting either *shc* (squalene hopene cyclase) or *hpnE* (hydroxysqualene oxidoreductase) resulted in a large growth deterioration at higher temperatures. It has previously been shown in other organisms that hopanoids are associated with sensitivity to high temperatures^9–12,31^, and MD simulations also suggest that hopanoids could reinforce membranes at higher temperatures^32^. Hopanoid biosynthesis deletion in both Δ*shc* and Δ*hpnE* mutants resulted in a large decrease in lipid packing measured *in vivo*, consistent with our previous observations with purified outer membranes^6^. Such low lipid packing, which is indicative of higher fluidity and lower mechanical robustness, could render the outer membrane susceptible to destabilization at higher temperatures, which can explain the growth impairment observed at higher temperatures. Interestingly, the change in lipid packing between 27 and 32°C was much higher for Δ*hpnE* and Δ*shc* mutant strains relative to the WT strain, suggesting impaired homeoviscous adaptation in the absence of hopanoids.

It has been hypothesized that carotenoids could share some of the lipid ordering properties of sterols^8,33,34^. Since Δ*hpnE* deletion eliminated both hopanoid and carotenoid synthesis, we had to target genes at a later stage of the carotenoid pathway that would allow to independently delete carotenoid synthesis to study the impact on growth and lipid packing. We targeted the genes involved in C_30_ biosynthesis; *crtN* and *crtP*, which both resulted in non-pigmented mutants that still produced hopanoids. Neither of the carotenoid mutants showed a significant growth impairment at any temperature from 10°C to 34°C which is a phenotype similar to what has been shown in *Acholeplasma*^35^, suggesting that in contrast to hopanoids, carotenoids are not critical for temperature adaptation. However, it is also possible that carotenoid deletion can be compensated for by other lipids, including hopanoids. Indeed, lipid packing increased in the non-pigmented mutants, indicating that carotenoids do have an influence on outer membrane properties, which agrees with what has been shown in *Pantoea sp*.^36^. Moreover, it has been shown that carotenoid production is increased at cold temperatures^15,16^. Carotenoids localize to the membrane bilayer and have a plethora of diverse structures which affect and localize in the membrane differently^37,38^. *In vitro* investigations have provided evidence that carotenoids have an ordering effect on membrane properties^8,33,39,40^. It had also been observed by ourselves and others^2,26,28^ that mutants deficient in hopanoid synthesis had much higher carotenoid content, indicating a possible crosstalk in the regulation of the two pathways. As carotenoids and hopanoids are both derived from squalene, it now seems likely that the increase in carotenoids which has been observed in Δ*shc* mutant strains could be due to an accumulation of squalene. Alternatively, carotenoids may serve in a different capacity unrelated to the physical properties of the membrane. For example, carotenoids play an important role in light scavenging in photosynthetic organisms^41^, and protecting cells from oxidative stress^36,42^. Nonetheless, our observations remain consistent with carotenoids playing a role in outer membrane physical homeostasis.

The presence of genes associated with both C_30_ and C_40_ carotenoid pathways, combined with evidence for the synthesis of only C_30_ carotenoids prodded us to examine the phylogeny of the two pathways for insights on their origins. An interesting clue emerged from the recent observation that squalene-derived carotenoids were also present in the hopanoid-producing Planctomycete, *Planctopirus limnophila*^26^. Why would such distantly related organisms possess both squalene-derived lipid synthesis pathways for carotenoids and hopanoids? We performed evolutionary analyses for the genes involved in the C_40_ and C_30_ carotenoid biosynthetic pathways and showed that they co-occur in the genomes of specific Alphaproteobacteria orders like Acetobacterales, Azospirillales and Rhodospirillales and Rhizobiales, the latter including *M. extorquens* (**Figure 5**). While the phylogeny of HpnCDE and C_40_ carotenoid enzymes suggest an ancestral feature of Alpha- and Gammaproteobacteria, the C_30_ carotenoid pathway in Alphaproteobacteria orders most likely represents a secondary acquisition by LGT from Planctomycetes. By confirming that *M. extorquens* produces only C_30_ carotenoids (**Figure 2, Figure S1**), and given that the enzyme CrtM is absent in Alphaproteobacteria we propose that the C_40_ carotenoid pathway via CrtB-CrtI-CrtD is not active for carotenoids production in *M. extorquens* at optimum growth conditions. Together, these observations suggest a replacement of C_40_ carotenoid biosynthesis in *M. extorquens* and possibly other related species. This replacement took advantage of the primitive squalene production via HpnCDE implying that the production of precursor for carotenoid and hopanoid biosynthesis in *M. extorquens* is controlled by the same genetic mechanisms. This fact also supports the notion that the two isoprenoid lipid classes (hopanoids and carotenoids) serve complementary roles in modulating membrane properties. We consequently hypothesize that the observed overexpression of carotenoids^2,18,26^ is due to squalene accumulation (**Table 1**) rather than it serving a compensatory role (**Figure 4**). The acquisition of the C_30_ pathway following the C_40_ pathway in *M. extorquens* would imply that C_30_ carotenoids impart an advantage, providing a new mystery to explore in *M. extorquens*.

*Methylobacterium extorquens* is on its way towards becoming a well-characterized and robust model system for studying the role of lipid structure in membrane function and organismal fitness. It was recently shown that *M. extorquens* has the simplest lipidome so far observed in any organism^17^, making it an ideal system for exploring the principles of lipidome adaptation. While the phospholipidome is relatively well-explored by comparison, the role of isoprenoid-lipids is still relatively undefined. Our observations raise the possibility that carotenoids and hopanoids serve complimentary roles in outer membrane adaptation. By revealing that carotenoids are squalene-derived and identifying genes in the carotenoid pathway, this study now provides a new tool to explore the property-function relationship of carotenoids and their relationship with hopanoids in *Methylobacterium*.

## Materials and Methods

### Media, Growth Conditions

*Methylobacterium* strains were grown at 30°C in minimal medium described by^43^ referred to as hypho medium, with 9.9 mM disodium succinate (Sigma Aldrich, W327700) as the carbon source at 160 rpm shaking (ISF1-X Kuhner shaker). *Escherichia coli* strains were grown at 37°C in LB medium (Carl Roth, X968). Triparental conjugation was performed on Nutrient broth medium (Carl Roth, X929.1). All solid media plates were prepared with 1.5% Agar-Agar (Carl Roth, 1347). Antibiotics for selection were at the following concentrations for *Methylobacterium*: Trimethoprim (Tmp) 10μg/ml (Cayman chemicals, 16473), Tetracycline (Tc) 10μg/ml (Carl Roth, HP63), Kanamycin 25μg/ml (Carl Roth, T832), for *E. coli*: Kanamycin (Km): 50μg/ml, Chloramphenicol (Cm) 25μg/ml (Sigma Aldrich, C1919). Plasmid pLC291^44^ was induced using Anhydrotetracycline hydrochloride 25ng/ml (Alfa Aesar, J66688).

### Evolutionary analyses for C_30_ and C_40_ carotenoid pathway

We performed protein searches of CrtI (P54980), CrtD (Q01671), CrtN (O07855) and CrtP (Q2FV57) against NCBI database using phmmer^45^ and with e-value threshold of 1e-5. We combined all the sequences obtained and using GTDB taxonomy^46^, we removed redundant sequences by taxonomic orders (from 90 up to 50% of identity threshold for the less and more represented groups respectively). We then aligned this set of non-redundant sequences using MAFFT^47^ and performed a fast phylogenetic tree using FastTree^48^ to exclude spurious sequences. Once we obtained the final set of sequences, we re-aligned with MAFFT and removed those enriched gap positions using trimAl^49^. For the final phylogenetic reconstruction, we used IQ-TREE^50^. We obtained branch supports with the ultrafast bootstrap^51^ and the evolutionary models were automatically selected using ModelFinder^52^ implemented in IQ-TREE and chosen according to BIC criterion.

For the phylogenetic profile, the distribution of HpnCDE, Sqs and CrtB enzymes were obtained from data previously generated^26^. The distribution of FAD-dependent desaturases was obtained from the phylogenetic reconstruction performed in this study. The taxonomic tree was obtained from GTDB repository (https://gtdb.ecogenomic.org/) pruning those sequences of interest. Phylogenetic trees were visualized and annotated in iTOL^53^.

### Growth rate at different temperatures

Fresh cells were passaged at least once in Erlenmeyer flasks at 30°C, cells were then diluted to OD_600_ of 0.02, and cultured in 96-half-deepwell microplate (enzyscreen, CR1469c). Cells were then grown at 650 rpm shaking on orbital thermoshaker (inheco, 7100146), at temperatures mentioned in the study. 10, 15, 20, 27, 30, 32, 34, 36°C. OD620 was measured on a plate reader (Molecular devices, FilterMax F3) included in an automated system (Beckman Coulter, Liquid Handler Biomek i7 Hybrid)

### Carotenoids extraction for absorbance scan

Bligh and Dyer extraction^54^ was used to extract carotenoids for cells grown to late exponential. 10 ml of cells of different *Methylobacterium* mutants were collected at 5000 rcf, for 10 minutes, washed once with 1x D-PBS. Wet weight of cell pellet was weighed. Cells were resuspended in water to 200μl (taking weight into account), adding 250μl chloroform (Carl Roth, Y015), and 500μl of methanol (VWR chemicals, 20903.368). The homogenous mixture was then sonicated in ultrasonic bath (Bandelin, Ultrasonic bath SONOREX DIGITEC DT 510 F) for 30 minutes.

Samples were centrifuged at 12000 rcf for 1 minute (Thermo Scientific, Microcentrifuge Pico™ 21), supernatant was collected and extracted by adding 250μl water, and 500μl chloroform, vigorously mixing the cells, and collecting the lower organic phase into a new tube. Extraction was repeated three times, then the collected extract was dried using vacuum concentrator (Christ, RVC 2-25 CD). Finally, the dried extract was dissolved in ethyl acetate (EtOAc) (Merck, 1.06923.2511) to a final concentration 0.1 mg/μl (of pellet wet weight). Absorbance Scan was performed on (Tecan, plate reader Spark M20) on the pigments in ethyl acetate, using cuvette (Hellma, 105-202-15-40).

### Isolation and Saponification of Carotenoids

Carotenoids extraction was adapted from^55^. Briefly, 10 mg wet weight of each sample was extracted using 1 ml methanol containing 6 % KOH and incubated for at least 14 h at 4° C in the dark. Supernatant was collected after centrifugation (1,500 g, 5 min) and reduced in a speedvac concentrator (Savant SPD111V; Thermo Fisher Scientific, Massachusetts, USA). EtOAc and saturated NaCl were added in equal volumes while thoroughly mixing after each addition. Upper organic phase was collected after centrifugation (10,000 g at 4°C for 5 min), washed twice with distilled water and completely dried.

### LC-MS Analysis

Dried extracts, squalene (Sigma Aldrich) and diplopterol (Chiron AS) were dissolved in acetonitrile and filtered with Minisart RC 4 (Sartorius, Stonehouse, UK) before applying 5 μl aliquots to a Acquity UPLC BEH C18 column (1.7 μm, 2.1 × 150 mm; Waters, Milford, Massachusetts) using an Agilent 1290 Infinity II HPLC system equipped with an diode array detector. Extracts were eluted with a gradient solvent system consisting of water with 0.1 % formic acid (A) and acetonitrile with 0.1 % formic acid (B). The gradient selected was: 0 min: 80 % B, 5 min: 80 % B, 15 min: 95 % B, 35 min: 95 % B, 40 min: 80 % B at a constant flow rate of 0.4 ml/min.

Carotenoids were identified using a combination of absorption spectra, retention time and mass spectra. For the differentiation between pigments with C_30_ and C_40_ carbon backbones Zeaxanthin and β-carotene (DHI Lab Products, Denmark) were run as examples for C_40_ pigments, and extracts from *Staphylococcus aurerus* 533 R4 (DSM 20231) and *Methylorubrum rhodinum* (DSM 2163) were prepared as controls for C_30_ pigments. Mass spectra were monitored in positive electron spray ionization (ESI) mode in a mass range of m/z 300 – 1000 on the Agilent 6545 Q-TOF system (Agilent, Waldbronn, Germany) using the following conditions: drying gas temperature 300 °C, drying gas flow rate 8 l/min, sheat gas temperature 350°C, sheat gas flow rate 12 l/min, capillary voltage 3000V.

### In vivo Di-4 spectroscopy

Three biological triplicates of cells were grown at either 27°C or 32°C until cultures reached mid exponential growth at around OD_600_ ~0.5. Cells were then diluted to OD_600_ 0.2, washed and resuspended in succinate-free media. Cells were then incubated with 80nM Di-4 ANEPPDHQ (ThermoFisher, D36802) for 10 minutes at 950 rpm shaking on a thermomixer (Eppendorf, ThermomixerC). Subsequently, cells were plated onto a black 96-well plate in analytical triplicates per sample, and measured in a plate reader (TecanSpark M20). Excitation was set to: 485 nm, and emission was recorded at 540 nm and 670 nm with a bandwidth of 20nm.

### Strains, Construction of Plasmids, Generation of Mutants, and Gene complementation

*Methylobacterium extorquens* PA1 with cellulose synthase deletion was used in this study and referred to hereafter as WT^43,44^, Δ*shc* was already available^6^. Genes for carotenoids biosynthesis were identified based on *M. extorquens* gene annotations for phytoene desaturase and phytoene synthases, and BLASTp was used to reconstruct the carotenoids biosynthesis pathway as shown Mutants were constructed by unmarked allelic exchange as described^56,57^, for each gene primers were designed to include 500bp upstream and 500bp downstream overhangs of the gene. The produced PCR product was then used as a template for the construction of 2 plasmids one to delete the gene and one for the inducible expression of the gene in the knockout strain as explained in (**Table 2**). For gene deletion: plasmid pCM433^56^ was linearized via restriction digestion using enzymes NotI-HF, and SacI-HF (NEB, R3189, R3156 respectively), overhangs upstream and downstream of the gene of interest were amplified (primers sequences available in **Table S1**) and purified then cloned into linearized pCM433 using In-Fusion HD Cloning plus kit (Takara), primer design was done using primer design tool (Takara). For inducible expression of the gene: plasmid pLC291 was linearized using EcoRI-Hf, and KpnI-HF restriction enzymes (NEB R3101, R3142 respectively), gene was then PCR amplified and purified then cloned into plasmid pLC291^44^ using In-Fusion HD Cloning plus kit (Takara). All PCR products and linearized vector were purified (Macherey and Nagel, Nucleospin PCR clean-up Gel extraction).

**Table 2.** Overhangs for gene deletion were introduced into pCM433^56^ Gene was cloned into pLC291 under the control of anhydrotetracycline inducible promoter^44^

| Gene | Protein ID | Deletion plasmid<sup>a</sup> | Inducible expression plasmid<sup>b</sup> |
| --- | --- | --- | --- |
| *hpnE* | WP_012253488.1 | pLMM013 | pLMM019 |
| *crtN* | WP_012254689.1 | pMG027 | pMG028 |
| *crtP* | WP_003603441.1 | pMG029 | pMG030 |
| *crtB* | WP_012254336.1 | pMG007 | — |

Deletion plasmids were introduced into WT via triparental conjugation. WT cells were mated with *E. Coli* pRK2073 helper cells, and E. coli Stellar cells that carry the deletion/expression plasmid, and the mating was done using a ratio of 5:1:1 Acceptor-strain : helper-strain : donor-strain. The conjugation was done on NB-Agar plates at 30°C, overnight, then the cells were plated on hypho media-agar plates with Tmp, and Tc. The clones were then grown for 9 hours in liquid media, then plated on 10% sucrose plates for selection of mutants. Colony PCR was then performed on clones from the sucrose plates, using Primers (**Table S1**) for gene template. PCR products of the correct size for gene deletion were then sequenced to confirm deletion of genes.

## Acknowledgments

The authors wish to thank members of the Sáenz group especially Grzegorz Chwastek, André Nadler, and Michael Schlierf for discussions; André Nadler for manuscript comments; Lisa-Maria Müller and Lisa Junghans for technical assistance. This work was supported by the B CUBE, TU Dresden, a German Federal Ministry of Education and Research BMBF grant (to J.S., project 03Z22EN12), and a VW Foundation ‘’Life’’ grant (to J.S., project 93090)

---

**Figure S1.**
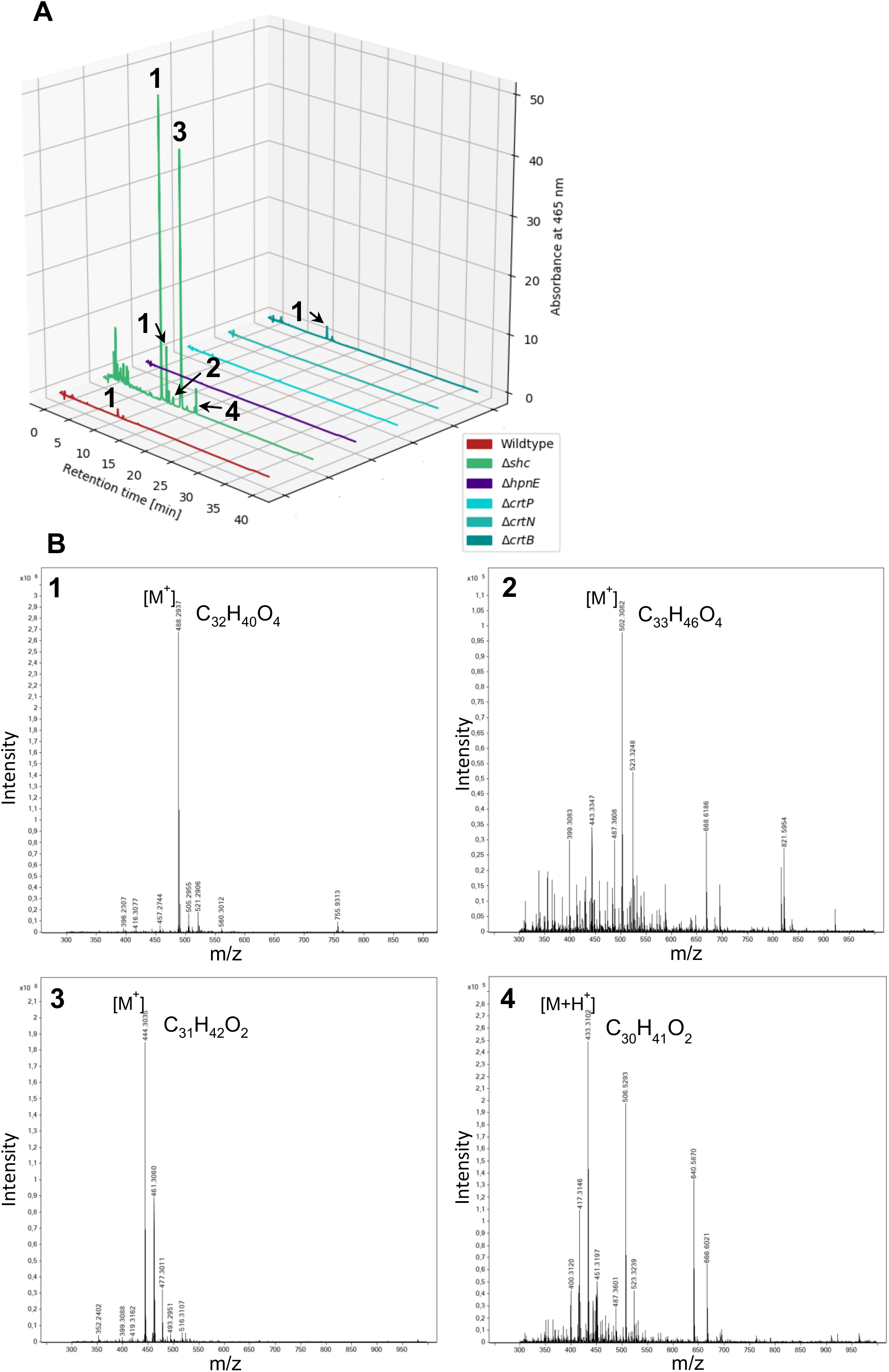
Identification of C_30_ carotenoids. **A**. Absorbance spectra at 465 nm of *M. extorquens* and mutant strains; Δ*shc*, Δ*hpnE*, Δ*crtP*, Δ*crtN* and Δ*crtB.* **B**. Mass spectra of the corresponding absorbance spectra peaks.

**Figure S2.**
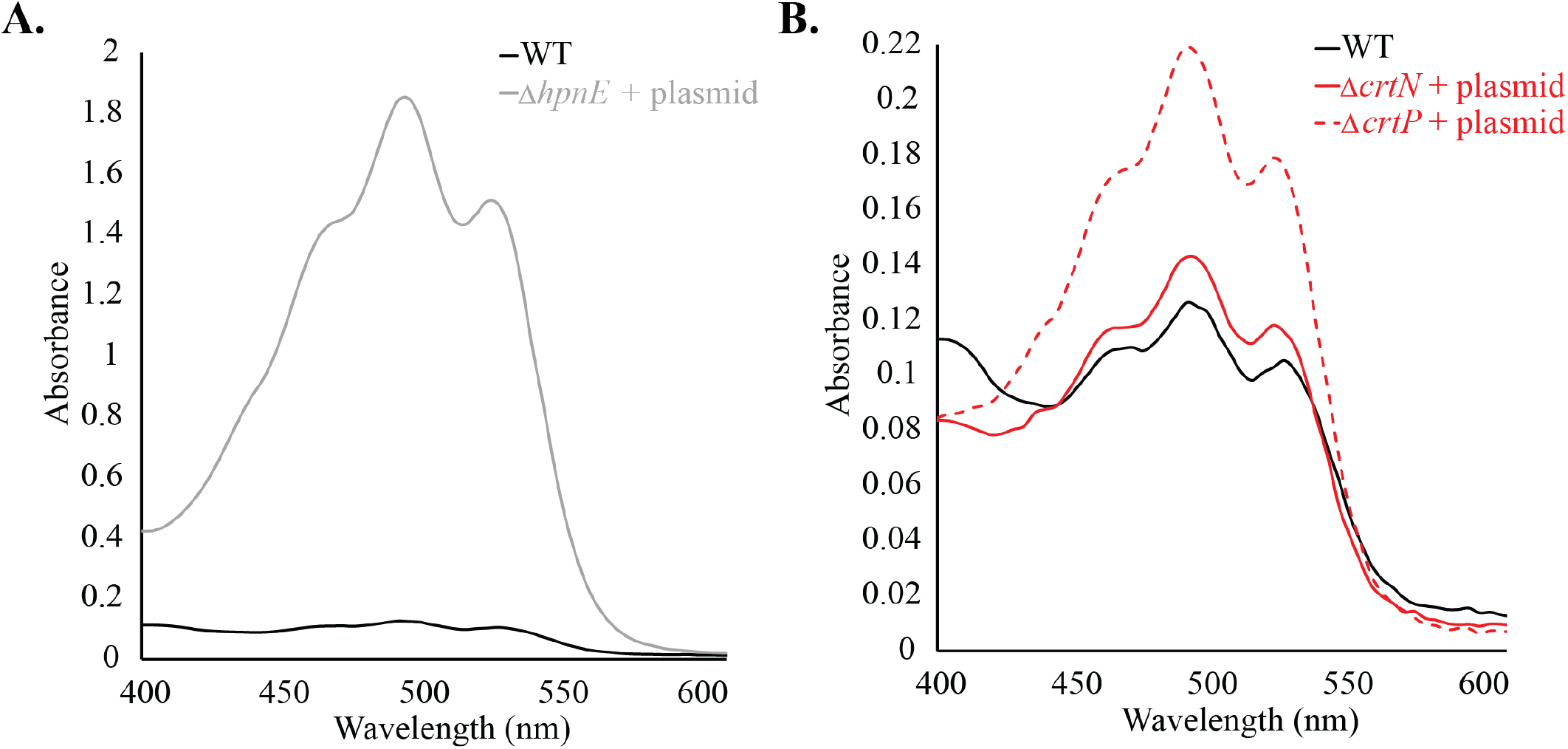
Absorbance spectra of lipids extracted from *M. extorquens* mutant strains upon expression of plasmids carrying the knocked-out genes **A.** Δ*hpnE* strain + plasmid pLMM019 **B.** Strains Δ*crtN*, and Δ *crtP* + plasmids pMG028, pMG030 respectively. (normalized to cell mass).

**Table S1:**
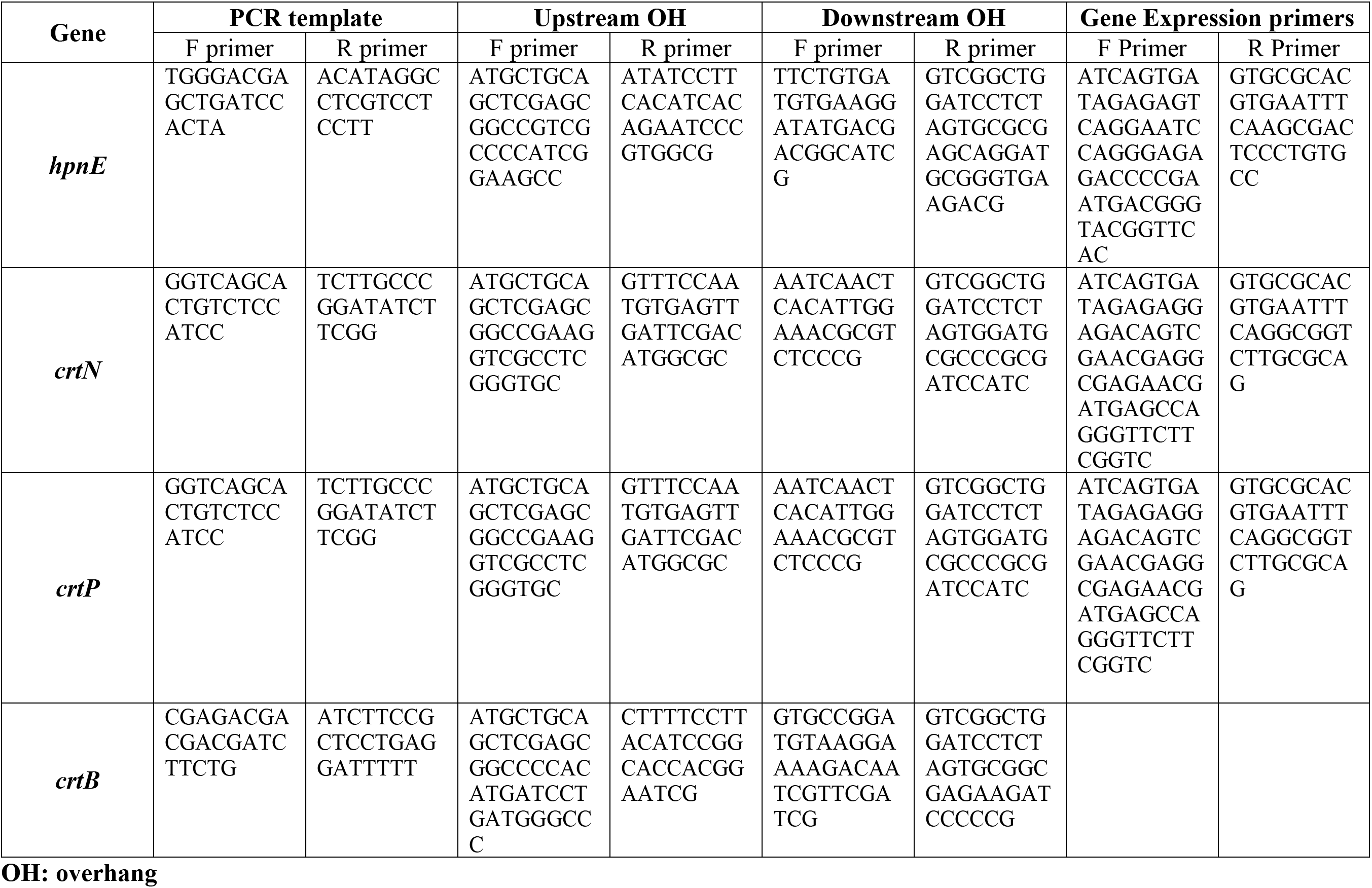

## References

1. Mouritsen, O. G. & Zuckermann, M. J. What’s so special about cholesterol? Lipids 39, 1101–1113 (2004).

2. Rivas-Marin, E. et al. Essentiality of sterol synthesis genes in the planctomycete bacterium Gemmata obscuriglobus. Nat Commun 10, 2916 (2019).

3. Ourisson, G., Rohmer, M. & Poralla, K. Prokaryotic Hopanoids and Other Polyterpenoid Sterol Surrogates. Ann. Rev. Microbiol. 41, 301–333 (1987).

4. Bramkamp, M. & Lopez, D. Exploring the Existence of Lipid Rafts in Bacteria. Microbiol Mol Biol R 79, 81–100 (2015).

5. Saenz, J. P., Sezgin, E., Schwille, P. & Simons, K. Functional convergence of hopanoids and sterols in membrane ordering. Proceedings of the National Academy of Sciences of the United States of America 1–5 (2012) doi:10.1073/pnas.1212141109/-/dcsupplemental/pnas.201212141si.pdf.

6. Saenz, J. P. et al. Hopanoids as functional analogues of cholesterol in bacterial membranes. Proceedings of the National Academy of Sciences of the United States of America 112, 11971–11976 (2015).

7. Silipo, A. et al. Covalently linked hopanoid-lipid A improves outer-membrane resistance of a Bradyrhizobium symbiont of legumes. Nat Commun 5, 5106 (2014).

8. Mostofian, B., Johnson, Q. R., Smith, J. C. & Cheng, X. Carotenoids promote lateral packing and condensation of lipid membranes. Phys Chem Chem Phys 22, 12281–12293 (2020).

9. Belin, B. J. et al. Hopanoid lipids: from membranes to plant–bacteria interactions. Nat Rev Microbiol 16, 304–315 (2018).

10. Doughty, D. M. et al. The RND-family transporter, HpnN, is required for hopanoid localization to the outer membrane of Rhodopseudomonas palustris TIE-1. Proc National Acad Sci 108, E1045–E1051 (2011).

11. Poralla, K., Härtner, T. & Kannenberg, E. Effect of temperature and pH on the hopanoid content of Bacillus acidocaldarius. FEMS Microbiology Letters 23, 2–3 (1984).

12. Schmidt, A., Bringer-Meyer, S., Poralla, K. & Sahm, H. Effect of alcohols and temperature on the hopanoid content of Zymomonas mobilis. Appl Microbiol Biot 25, 32–36 (1986).

13. Kulkarni, G., Wu, C.-H. & Newman, D. K. The General Stress Response Factor EcfG Regulates Expression of the C-2 Hopanoid Methylase HpnP in Rhodopseudomonas palustris TIE-1. J Bacteriol 195, 2490–2498 (2013).

14. Chattopadhyay, M. K. & Jagannadham, M. V. Maintenance of membrane fluidity in Antarctic bacteria. Polar Biology 24, 386–388 (2001).

15. Fong, N., Burgess, M., Barrow, K. & Glenn, D. Carotenoid accumulation in the psychrotrophic bacterium Arthrobacter agilis in response to thermal and salt stress. Appl Microbiol Biot 56, 750–756 (2001).

16. Seel, W. et al. Carotenoids are used as regulators for membrane fluidity by Staphylococcus xylosus. Sci Rep-uk 10, 330 (2020).

17. Chwastek, G. et al. Principles of Membrane Adaptation Revealed through Environmentally Induced Bacterial Lipidome Remodeling. Cell Reports 32, 108165 (2020).

18. Bradley, A. S. et al. Hopanoid-free Methylobacterium extorquens DM4 overproduces carotenoids and has widespread growth impairment. Plos One 12, e0173323 (2017).

19. Pan, J.-J. et al. Biosynthesis of Squalene from Farnesyl Diphosphate in Bacteria: Three Steps Catalyzed by Three Enzymes. Acs Central Sci 1, 77–82 (2015).

20. Furubayashi, M., Li, L., Katabami, A., Saito, K. & Umeno, D. Construction of carotenoid biosynthetic pathways using squalene synthase. Febs Lett 588, 436–442 (2014).

21. Amaro, M., Reina, F., Hof, M., Eggeling, C. & Sezgin, E. Laurdan and Di-4-ANEPPDHQ probe different properties of the membrane. J Phys D Appl Phys 50, 134004 (2017).

22. Ma, Y., Benda, A., Kwiatek, J., Owen, D. M. & Gaus, K. Time-Resolved Laurdan Fluorescence Reveals Insights into Membrane Viscosity and Hydration Levels. Biophys J 115, 1498–1508 (2018).

23. Steinkühler, J., Sezgin, E., Urbančič, I., Eggeling, C. & Dimova, R. Mechanical properties of plasma membrane vesicles correlate with lipid order, viscosity and cell density. Commun Biology 2, 337 (2019).

24. Lorent, J. H. et al. Plasma membranes are asymmetric in lipid unsaturation, packing and protein shape. Nat Chem Biol 16, 644–652 (2020).

25. Dien, S. J. V., Marx, C. J., O’Brien, B. N. & Lidstrom, M. E. Genetic characterization of the carotenoid biosynthetic pathway in Methylobacterium extorquens AM1 and isolation of a colorless mutant. Applied and Environmental Microbiology 69, 7563–7566 (2003).

26. Santana-Molina, C., Rivas-Marin, E., Rojas, A. M. & Devos, D. P. Origin and evolution of polycyclic triterpene synthesis. Mol Biol Evol 37, 1925–1941 (2020).

27. Bradley, A. S., Pearson, A., Saenz, J. P. & Marx, C. J. Adenosylhopane: The first intermediate in hopanoid side chain biosynthesis. Organic Geochemistry 41, 1075–1081 (2010).

28. Bradley, A. S. et al. Hopanoid-free Methylobacterium extorquens DM4 overproduces carotenoids and has widespread growth impairment. PLoS ONE 12, e0173323–18 (2017).

29. Csáky, Z. et al. Squalene lipotoxicity in a lipid droplet-less yeast mutant is linked to plasma membrane dysfunction. Yeast 37, 45–62 (2020).

30. Valachovic, M., Garaiova, M., Holic, R. & Hapala, I. Squalene is lipotoxic to yeast cells defective in lipid droplet biogenesis. Biochem Bioph Res Co 469, 1123–1128 (2016).

31. Kulkarni, G., Wu, C.-H. & Newman, D. K. The general stress response factor EcfG regulates expression of the C-2 hopanoid methylase HpnP in Rhodopseudomonas palustris TIE-1. Journal of Bacteriology 195, 2490–2498 (2013).

32. Caron, B., Mark, A. E. & Poger, D. Some Like It Hot: The Effect of Sterols and Hopanoids on Lipid Ordering at High Temperature. J Phys Chem Lett 5, 3953–3957 (2014).

33. Subczynski, W. K., Markowska, E., Gruszecki, W. I. & Sielewiesiuk, J. Effects of polar carotenoids on dimyristoylphosphatidylcholine membranes: a spin-label study. Biochimica Et Biophysica Acta Bba - Biomembr 1105, 97–108 (1992).

34. Socaciu, C., Jessel, R. & Diehl, H. A. Competitive carotenoid and cholesterol incorporation into liposomes: effects on membrane phase transition, fluidity, polarity and anisotropy. Chem Phys Lipids 106, 79–88 (2000).

35. Razin, S. & Rottem, S. Role of carotenoids and cholesterol in the growth of Mycoplasma laidlawii. J Bacteriol 93, 1181–1182 (1967).

36. Kumar, S. V. et al. Loss of carotenoids from membranes of Pantoea sp. YR343 results in altered lipid composition and changes in membrane biophysical properties. Biochimica Et Biophysica Acta Bba - Biomembr 1861, 1338–1345 (2019).

37. Wisniewska, A., Widomska, J. & Subszynski, W. K. Carotenoid-membrane interactions in liposomes: Effect of dipolar, monopolar, and nonpolar carotenoids. Acta Biochimica Polonica 53, 475–484 (2006).

38. Milon, A., Wolff, G., Ourisson, G. & Nakatani, Y. Organization of Carotenoid-Phospholipid Bilayer Systems. Incorporation of Zeaxanthin, Astaxanthin, and their C50 Homologues into Dimyristoylphosphatidylcholine Vesicles. Helv Chim Acta 69, 12–24 (1986).

39. Kostecka-Gugała, A., Latowski, D. & Strzałka, K. Thermotropic phase behaviour of α-dipalmitoylphosphatidylcholine multibilayers is influenced to various extents by carotenoids containing different structural features-evidence from differential scanning calorimetry. Biochimica Et Biophysica Acta Bba - Biomembr 1609, 193–202 (2003).

40. Gabrielska, J. & Gruszecki, W. I. Zeaxanthin (dihydroxy-β-carotene) but not β-carotene rigidities lipid membranes: A 1H-NMR study of carotenoid-egg phosphatidylcholine liposomes. Biochimica et Biophysica Acta - Biomembranes 1285, 167–174 (1996).

41. Polívka, T. & Frank, H. A. Molecular Factors Controlling Photosynthetic Light Harvesting by Carotenoids. Accounts Chem Res 43, 1125–1134 (2010).

42. Kim, M., Seo, D.-H., Park, Y.-S., Cha, I.-T. & Seo, M.-J. Isolation of Lactobacillus plantarum subsp. Plantarum Producing C_30_ Carotenoid 4,4’-Diaponeurosporene and the Assessment of Its Antioxidant Activity. J Microbiol Biotechn 29, 1925–1930 (2019).

43. Delaney, N. F. et al. Development of an Optimized Medium, Strain and High-Throughput Culturing Methods for Methylobacterium extorquens. PLoS ONE 8, e62957–17 (2013).

44. Chubiz, L. M., Purswani, J., Carroll, S. M. & Marx, C. J. A novel pair of inducible expression vectors for use in Methylobacterium extorquens. BMC Research Notes 6, 1–8 (2013).

45. Finn, R. D., Clements, J. & Eddy, S. R. HMMER web server: interactive sequence similarity searching. Nucleic Acids Res 39, W29–W37 (2011).

46. Parks, D. H. et al. A standardized bacterial taxonomy based on genome phylogeny substantially revises the tree of life. Nat Biotechnol 36, 996–1004 (2018).

47. Katoh, K. & Standley, D. M. MAFFT Multiple Sequence Alignment Software Version 7: Improvements in Performance and Usability. Mol Biol Evol 30, 772–780 (2013).

48. Price, M. N., Dehal, P. S. & Arkin, A. P. FastTree: Computing Large Minimum Evolution Trees with Profiles instead of a Distance Matrix. Mol Biol Evol 26, 1641–1650 (2009).

49. Capella-Gutiérrez, S., Silla-Martínez, J. M. & Gabaldón, T. trimAl: a tool for automated alignment trimming in large-scale phylogenetic analyses. Bioinformatics 25, 1972–1973 (2009).

50. Nguyen, L.-T., Schmidt, H. A., Haeseler, A. von & Minh, B. Q. IQ-TREE: A Fast and Effective Stochastic Algorithm for Estimating Maximum-Likelihood Phylogenies. Mol Biol Evol 32, 268–274 (2015).

51. Hoang, D. T., Chernomor, O., Haeseler, A. von, Minh, B. Q. & Vinh, L. S. UFBoot2: Improving the Ultrafast Bootstrap Approximation Brief Communication Open Access. MBE (2017) doi:https://doi.org/10.1093/molbev/msx281.

52. Kalyaanamoorthy, S., Minh, B. Q., Wong, T. K. F., Haeseler, A. von & Jermiin, L. S. ModelFinder: fast model selection for accurate phylogenetic estimates. Nat Methods 14, 587–589 (2017).

53. Letunic, I. & Bork, P. Interactive Tree Of Life (iTOL) v4: recent updates and new developments. Nucleic Acids Res 47, gkz239- (2019).

54. Bligh, E. G. & Dyer, W. J. A Rapid Method of Total Lipid Extraction and Purification. Canadian Journal of Biochemistry and Physiology 37, (1959).

55. Kim, S. H. & Lee, P. C. Functional Expression and Extension of Staphylococcal Staphyloxanthin Biosynthetic Pathway in Escherichia coli. J Biol Chem 287, 21575–21583 (2012).

56. Marx, C. J. Development of a broad-host-range sacB-based vector for unmarked allelic exchange. BMC Research Notes 1, 1–8 (2008).

57. Hmelo, L. R. et al. Precision-engineering the Pseudomonas aeruginosa genome with two-step allelic exchange. Nat Protoc 10, 1820–1841 (2015).

